# Low intensity repetitive transcranial magnetic stimulation modulates brain-wide functional connectivity to promote anti-correlated activity

**DOI:** 10.1101/2022.08.13.503840

**Authors:** Jessica Moretti, Dylan J. Terstege, Eugenia Z. Poh, Jonathan R. Epp, Jennifer Rodger

**Author notes:** **Corresponding Author:** Jessica Moretti;, Jennifer Rodger;, M317, School of Biological Sciences, The University of Western Australia, 35 Stirling Highway, Crawley WA 6009. These two authors contributed equally (Jessica Moretti, Dylan J. Terstege). Netherlands Institute for Neuroscience, Amsterdam, The Netherlands (Eugenia Z. Poh).

## Abstract

**Background:** Repetitive transcranial magnetic stimulation (rTMS) induces action potentials to induce plastic changes in the brain with increasing evidence for the therapeutic importance of brain-wide functional network effects of rTMS; however, the influence of sub-action potential threshold (low-intensity; LI-) rTMS on neuronal activity is largely unknown.

**Hypothesis:** We investigated whether LI-rTMS modulates neuronal activity and functional connectivity. We also specifically assessed modulation of parvalbumin interneuron activity.

**Methods:** We conducted a brain-wide analysis of c-Fos, a marker for neuronal activity, in mice that received LI-rTMS to visual cortex. Mice received single or multiple sessions of excitatory 10Hz LI-rTMS with custom rodent coils or were sham controls. We assessed changes to c-Fos positive cell densities and c-Fos/parvalbumin co-expression. Peak c-Fos expression corresponded with activity during rTMS. We also assessed functional connectivity changes using brain-wide c-Fos-based network analysis.

**Results:** LI-rTMS modulated c-Fos expression in cortical and subcortical regions. c-Fos density changes were most prevalent with acute stimulation, however chronic stimulation decreased parvalbumin interneuron activity, most prominently in the amygdala and striatum. LI-rTMS also increased anti-correlated functional connectivity, with the most prominent effects also in the amygdala and striatum following chronic stimulation.

**Conclusion:** LI-rTMS induces changes in c-Fos expression that suggest modulation of neuronal activity and functional connectivity throughout the brain. Our results suggest that LI-rTMS promotes anticorrelated functional connectivity, possibly due to decreased parvalbumin interneuron activation induced by chronic stimulation. These changes may underpin therapeutic rTMS effects, therefore modulation of subcortical activity supports rTMS for treatment of disorders involving subcortical dysregulation.

**Highlights:**

- Low-intensity rTMS increases brain-wide anti-correlated functional connectivity
- Acute excitatory LI-rTMS modulates cortical and subcortical neuronal activity
- Decreased parvalbumin interneuron activity may promote anti-correlated activity
- Striatum and amygdala show prominent modulation with LI-rTMS

## Introduction

Use of repetitive transcranial magnetic stimulation (rTMS) is increasing for the treatment of a range of neurological conditions, however there is still limited understanding of the effects of electromagnetic stimulation in the brain. Conventional rTMS is generally linked to direct electromagnetic activation of cortical tissue underneath the coil, where induced electric fields lead to plastic changes in the brain. However, rTMS-induced changes in brain activity can also occur outside of the initial stimulation site, which is thought to be due to indirect modulation of connected brain structures [1–4]; these significant changes to network connectivity may underpin the therapeutic effects of rTMS [5–11]. As a result, there has been growing interest in understanding the brain-wide changes in functional connectivity in response to rTMS. However, in humans, understanding this relationship is limited to EEG and fMRI techniques, which have limitations in spatial and temporal resolution, and due to technical restrictions of ferromagnetic TMS coils, are difficult to apply during stimulation. BOLD changes are also indirectly related to neural activity and therefore are unable to separate the contribution of different neuronal subtypes [12].

In order to explore rTMS-induced changes in neuronal activity at cellular resolution (without electrophysiology), previous studies used the immediate-early gene c-Fos as an indirect marker of neuronal activity in the brains of mice that had received rTMS [13–15]. However, those studies used a human rTMS coil, which was too large to deliver focal stimulation to the small mouse brain, precluding the study of connectivity changes [16]. To better emulate the spatial characteristics of human rTMS, and provide the opportunity to study activation of networks downstream of defined brain regions, here we delivered stimulation to the mouse brain using a miniaturised coil [17]. Despite the low intensity magnetic field delivered by these miniaturised coils (low-intensity (LI-) rTMS), they have been shown to induce a range of neurobiological changes in rodents, including changes in resting state connectivity that are comparable to those observed in humans [1,2,18]. A further advantage of the miniaturised coils is that they can be attached to the head of awake and freely moving animals, avoiding the confounds of restraint and anaesthesia required by larger coils [19,20].

To better understand how rTMS alters activity in, and connectivity between, different parts of the brain, we conducted a brain-wide analysis of c-Fos positive (c-Fos^+^) cell density in mice that were euthanised 90 minutes after LI-rTMS over the visual cortex to capture the peak of c-Fos expression corresponding with brain activity during stimulation. We specifically included analysis of c-Fos^+^ parvalbumin positive (PV^+^) neurons, which are usually GABA-ergic interneurons and play an important role in coordinating and modulating neuronal circuit activity [21,22], particularly in response to rTMS [15,23–26]. We then conducted a network analysis, correlating c-fos expression between brain regions to explore how functional connectivity changes during rTMS, and whether PV^+^ neurons may contribute to these changes [27–30]. Additionally, although therapeutic rTMS effects are often thought to be cumulative, there is still limited understanding of the different outcomes on brain activity of single and multiple sessions of rTMS [31, although see 32]. Therefore, we included animals that received either acute (single session) or chronic (14 daily sessions) of rTMS to visualise on a cellular level whether acute and chronic stimulation activate different brain regions and circuits.

## Methods

### Animals

All experiments were approved by The University of Western Australia Animal Ethics Committee (AEC 100/1639). Twenty-two wildtype C57Bl6/J (8wks) were used and housed in 12-h day/night cycle (7am-7pm). Mice were organised into acute or chronic stimulation groups, and either sham or LI-rTMS conditions (Acute Sham: *n =* 6, 3 males; Acute LI-rTMS: *n =* 6, 3 males; Chronic Sham: *n =* 5, 3 males; Chronic LI-rTMS: *n* =5, 2 males).

### Procedure

#### Coil Support Attachment Surgery

To allow stimulation to be delivered accurately on freely moving animals, surgery to attach a coil support to the skull was performed, as described previously [33]. The coil support consisted of a plastic pipette tip attached to the exposed skull over the stimulation target area with dental cement and trimmed to <10 mm in length. The skin was then sutured over the cement base of the coil support. Post-operatively, mice were housed individually with the cage hopper removed to prevent damage to the support, with Hydrogel (HydroGel, ClearH2O) and food provided *ad libitum*.

#### Stimulation

The coil support allows a custom LI-rTMS coil to be fixed in place during stimulation by attaching it to the support with an alligator clip. From the fifth day following surgery, mice were habituated to the coil by attaching a dummy coil to the support for 5-10 minutes each day for 3 days prior to beginning stimulation. Stimulation was delivered with a custom animal LI-rTMS coil (300 copper windings, external diameter, 8 mm; internal diameter 5 mm) delivering approximately 21 mT at the base of the coil. The coil was powered by an electromagnetic pulse generator (e-cell™) programmed to deliver 10Hz stimulation for 10 minutes (6000 pulses). Stimulation was applied to freely moving animals in their home cage either once (acute group), or daily for 14 days (chronic). For sham stimulation, the coil was attached to the support, but with the pulse generator switched off.

#### Tissue Processing

##### Tissue Collection

Animals were euthanised with sodium pentobarbitone (0.1ml i.p., Lethabarb, Virbac, Australia) on the final day of stimulation, 90 minutes after the beginning of stimulation to capture the peak c-Fos expression during stimulation. Animals were then transcardially perfused with saline (0.9 % NaCl, w/v) and paraformaldehyde (4% in 0.1 M phosphate buffer, w/v), the brains were dissected out and post-fixed in paraformaldehyde for 24h and transferred to 30% sucrose in phosphate buffer solution (PBS) (w/v) for cryoprotection. Coronal sections (30 μm) were cryosected into 5 series. One of the resulting series was divided in half, wherein sections were alternatingly sorted for either brain-wide c-Fos labelling or an analysis of parvalbumin and c-Fos co-expression. This division resulted in a spacing of 300μm between sections in each immunohistochemistry procedure.

##### Immunohistochemistry

###### Brain-Wide c-Fos Expression

Tissue sections were stained with c-Fos (Rabbit polyclonal c-Fos antibody, 1:5000, Abcam, ab190289) and NeuN (mouse monoclonal anti-NeuN, 1:2000, Millipore, MAB377). Free-floating sections were washed (30 minutes per wash) with PBS and permeabilised with two washes of 0.1 % Triton-X in PBS (PBS-T). Sections were incubated for 2h in blocking buffer of 3% bovine serum albumin (BSA, Sigma) and 2% donkey serum (Sigma) diluted in PBS-T. Primary antibodies were incubated in fresh blocking buffer at 4 °C for 18 hours, washed with PBS-T and then incubated with secondary antibodies for 2h (donkey anti-rabbit lgG Alexa Fluor 488, Invitrogen, Thermo Fisher, A21206; donkey anti-Mouse lgG Alexa Fluor 555, Invitrogen, Thermo Fisher, A21202, 1:600 in blocking buffer). Sections were washed twice with PBS before being mounted onto gelatin subbed slides, coverslipped with mounting medium (Dako, Glostrup, Denmark) and sealed with nail polish. Slides were stored at 4 °C in a light-controlled environment until imaging.

###### Parvalbumin and c-Fos Co-Expression

Tissue sections were washed three times (10 minutes per wash) in 0.1M PBS before being incubated in a primary antibody solution of mouse anti-PV (1:2000, EnCor Biotechnology Inc., MCA-3C9), rabbit anti-c-Fos (1:2000, EnCor Biotechnology Inc., RPCA-c-Fos), 3% normal goat serum, and 0.03% Triton-X100 for 48h. Tissue sections were washed three more times in 0.1M PBS before secondary antibody incubation. The secondary antibody solution was composed of 1:500 goat anti-mouse Alexa Fluor 594 (Invitrogen, Thermo Fisher, A11005) and 1:500 goat anti-rabbit Alexa Fluor 647 (Jackson ImmunoResearch, 111-605-003) in PBS for 24h. Sections were then transferred to 1:1000 DAPI solution for 20 minutes before three final PBS washes. Labelled sections were mounted to plain glass slides and coverslipped with PVA-DABCO mounting medium.

###### Imaging

For the analysis of brain-wide c-Fos expression density, tissue sections were imaged using a Nikon C2 Confocal microscope (Nikon, Tokyo, Japan). The entire section was imaged *via* multiple images taken at 10x magnification and z-stacks separated by 5μm. Images were automatically stitched together with a 10% overlap using NIS Elements software (Nikon, Tokyo, Japan).

Images of c-Fos and parvalbumin co-expression were collected using an OLYMPUS VS120-L100-W slide-scanning microscope (Richmond Hill, ON, Canada). Images of a single z-plane were collected using a 10x objective.

###### Image Processing

Quantification of c-Fos labelling was segmented and registered using a semi-automated pipeline described in Terstege et al. [34]. Briefly, c-Fos labelled cells were segmented using *Ilastik*, a machine-learning based pixel classification program [35]. *Ilastik* output images were then registered to the Allen Mouse Brain Atlas using *Whole Brain*, an R based software [36] and used in combination with custom *ImageJ* software designed to calculate region volumes and output accurate c-Fos density counts per region.

For analyses of c-Fos and parvalbumin co-expression, cells expressing c-Fos and parvalbumin labels were segmented independently using *Ilastik*. The *Ilastik* binary object prediction images were further processed using a custom *ImageJ* plug-in to identify instances of co-expression and output a binary image containing only these c-Fos and parvalbumin co-expressing cells. Finally, both these co-expression images and the *Ilastik* object prediction images of parvalbumin labelling were mapped to a custom neuroanatomical atlas based on a higher-order region organization of the Allen Mouse Brain Atlas using *FASTMAP* [37]. This approach facilitated the accurate assessment of the percentage of parvalbumin interneurons which were expressing c-Fos across several higher-order brain regions.

### Data Analysis

#### Brain-Wide c-Fos Density

To assess general activation of regions across the brain, c-Fos^+^ cells were quantified in 115 neuroanatomical regions. This regional organization encompassed the entire mouse brain and was selected based on experimenter ability to delineate these neuroanatomical regions of interest in NeuN-stained tissue (see Supplementary File S1 for list of regions and abbreviations).

We compared c-Fos expression density (c-Fos^+^ cells/mm^2^) in a negative binomial generalised linear model with a log link for each region of interest. Fixed factors were Stimulation and Time. Values with a Cook’s Distance > 0.5 were excluded and regions with less than three values in any group were excluded from analysis, resulting in 73 regions analysed for density. To account for multiple comparisons across regions we used a false discovery rate approach (Q = 0.01) for the omnibus effects. For regions that had significant omnibus effects, we followed up with analysis of the main effects and interaction. If there was a significant interaction effect, we ran simple main effect analyses in order to interpret the changes.

#### Functional Connectivity Networks

The impact of regional changes in c-Fos expression density on brain dynamics was examined through the scope of functional connectivity networks [27–30]. Networks were constructed by cross-correlating regional c-Fos expression density within each group to generate pairwise correlation matrices. Correlations were filtered by statistical significance (α < 0.005) and a false discovery rate of 95%. The number of pairwise correlations exhibiting anticorrelated activity and the mean Pearson’s correlation coefficient were assessed for each network. Network density, defined as the proportion of actual functional connections relative to the potential number of connections in a fully saturated network was also assessed [38].

#### Brain-Wide Parvalbumin and c-Fos Co-Expression

Regional co-expression of c-Fos and parvalbumin was expressed as a percentage of the total number of parvalbumin interneurons present in each region. These data were compared separately for acute and chronic groups using Two-Factor ANOVA, with factors of Stimulation and Region.

### Data Availability

All datasets generated for this study and the scripts developed for its analysis can be found at the following GitHub repository [https://github.com/dterstege/PublicationRepo/tree/main/Moretti2022].

## Results

### Brain Wide c-Fos Density Changes

LI-rTMS modulated c-Fos expression in various regions throughout the brain. Of the 73 regions included in analysis, 53 had a significant omnibus model effect. Several regions showed a significant effect of time, indicating differences between the chronic and acute group, regardless of stimulation. c-Fos density for regions showing significant changes are reported in Figures 1 and 2, organised by hierarchical brain region. A summary of significant effects and interaction are reported in Supplementary File S2. Percentage changes in c-Fos density between sham and LI-rTMS groups for all regions are reported in Supplementary Figure S1.

**Figure 1.**
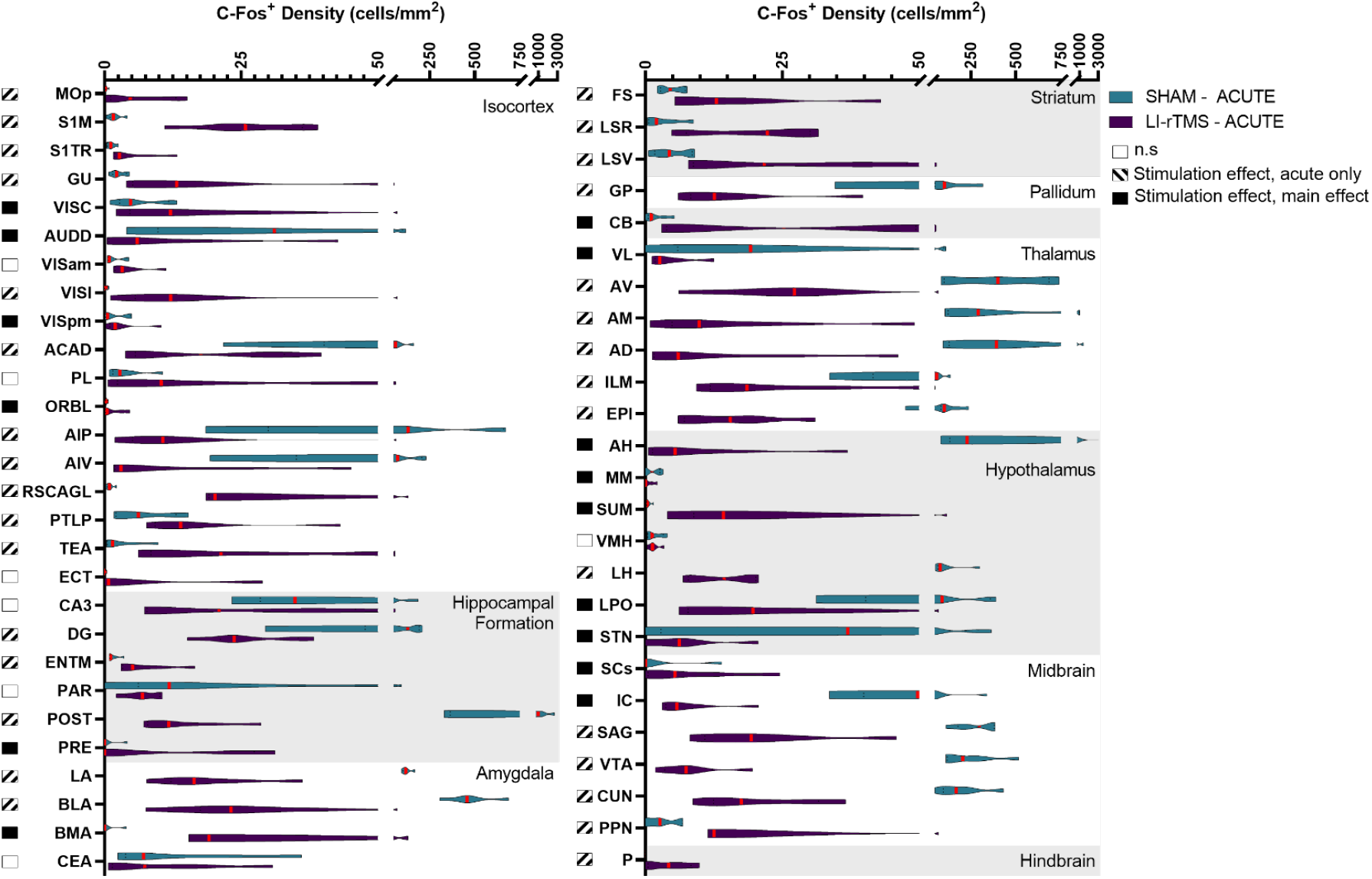
C-Fos cell density in mice in the acute stimulation group for regions which had a significant effect related to stimulation across all groups. Violin graphs represent c-Fos cell density (cells/mm^2^) for mice that received acute active or sham stimulation organised by hierarchical brain regions. Red lines indicate the median value. Shaded boxes to the left of the region name indicate whether there was a significant stimulation effect for the region.

**Figure 2.**
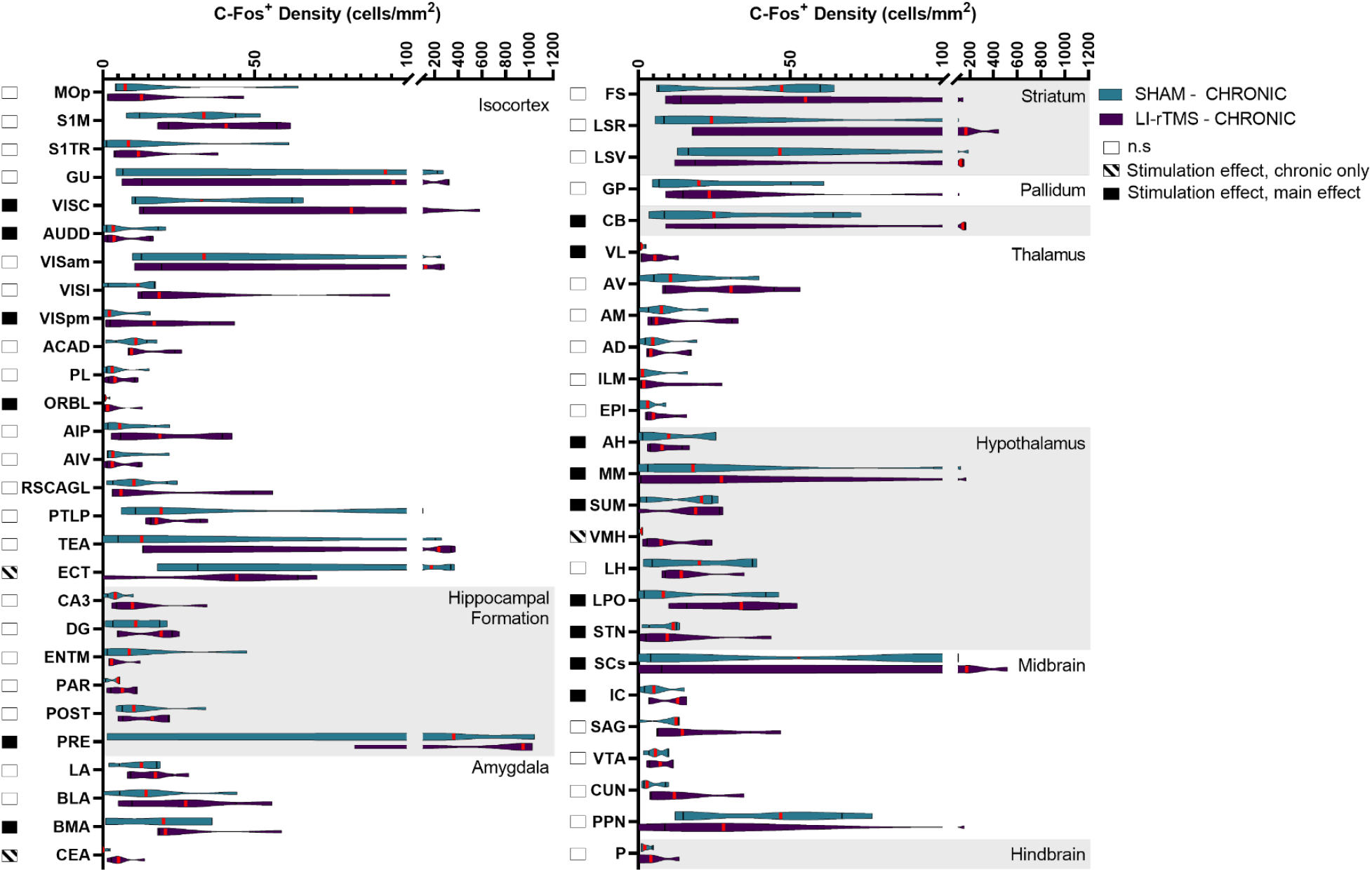
C-Fos cell density in mice in the chronic stimulation group for regions which had a significant effect related to stimulation across all groups. Violin graphs represent c-Fos cell density (cells/mm^2^) for mice that received acute active or sham stimulation organised by hierarchical brain regions. Red lines indicate the median value. Shaded boxes to the left of the region name indicate whether there was a significant stimulation effect for the region.

Stimulation-induced changes in c-Fos density were present throughout both cortical and subcortical regions - 49 regions showed an effect related to stimulation, and several had significant stimulation*time interactions. Follow up simple main effect analysis indicated that the main effect of stimulation could be interpreted for 14 regions, but the majority of stimulation-induced changes were due to altered activity following acute, but not chronic stimulation. Only 3 regions (ECT, CEA, VMH) showed significantly different c-Fos density following chronic but not acute stimulation. The direction of c-Fos density changes was varied across brain regions (24 regions show reduced c-Fos density; 24 regions show increased density). The direction of change in c-Fos expression for broader hierarchical groupings, based on significant changes in individual regions, is reported in Table 1.

**Table 1.**
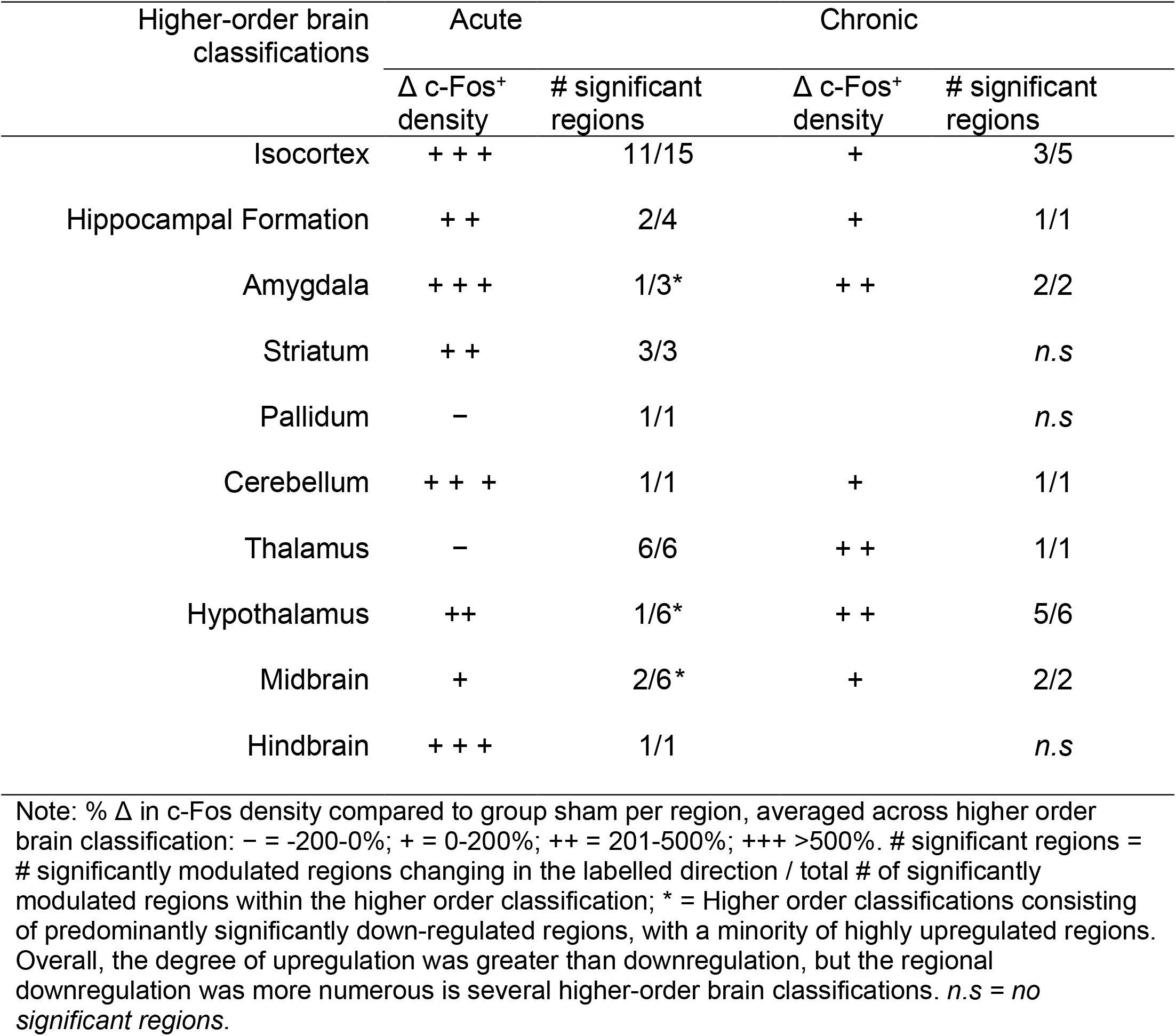
General direction of change in c-Fos expression following Acute or Chronic LI-rTMS, organised across higher order brain classifications.

| Higher-order brain classifications | Acute |  | Chronic |  |
| --- | --- | --- | --- | --- |
| | $\Delta$ c-Fos<sup>+</sup> density | # significant regions | $\Delta$ c-Fos<sup>+</sup> density | # significant regions |
| Isocortex | +++ | 11/15 | + | 3/5 |
| Hippocampal Formation | ++ | 2/4 | + | 1/1 |
| Amygdala | +++ | 1/3* | ++ | 2/2 |
| Striatum | ++ | 3/3 |  | *n.s* |
| Pallidum | – | 1/1 |  | *n.s* |
| Cerebellum | +++ | 1/1 | + | 1/1 |
| Thalamus | – | 6/6 | ++ | 1/1 |
| Hypothalamus | ++ | 1/6* | ++ | 5/6 |
| Midbrain | + | 2/6* | + | 2/2 |
| Hindbrain | +++ | 1/1 |  | *n.s* |

Note: % $\Delta$ in c-Fos density compared to group sham per region, averaged across higher order brain classification: – = -200-0%; + = 0-200%; ++ = 201-500%; +++ >500%. # significant regions = # significantly modulated regions changing in the labelled direction / total # of significantly modulated regions within the higher order classification; \* = Higher order classifications consisting of predominantly significantly down-regulated regions, with a minority of highly upregulated regions. Overall, the degree of upregulation was greater than downregulation, but the regional downregulation was more numerous in several higher-order brain classifications. *n.s* = *no significant regions*.

The largest difference in c-Fos expression was upregulation occurring during acute stimulation. Areas with the largest mean difference in c-Fos density with acute stimulation were prevalent in the cortex, as well as striatal regions. Downregulation was particularly prevalent in several thalamic regions during acute stimulation. In relation to the position of the coil, superficial regions positioned below the greatest induced e-field [17] showed significant increases in c-Fos with acute LI-rTMS (Fig. 3). Videos showing the 3D model of significantly regulated regions can be found in supplementary materials (Movie S1-S2), and the 3D objects from the videos, created using the Scalable Brain Atlas [39,40] are available in our the GitHub repository.

**Figure 3.**
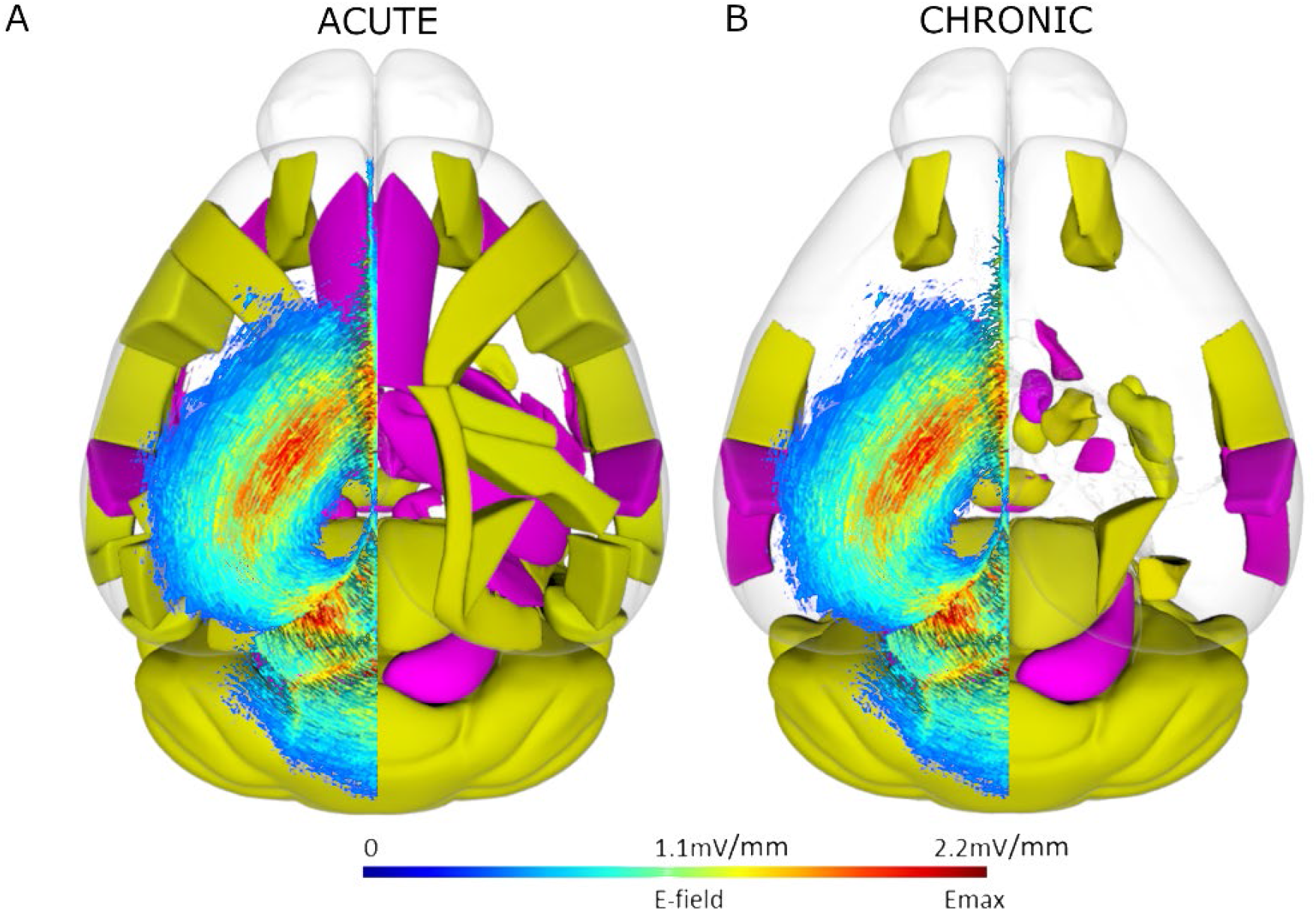
Spatial representation superficial brain regions with significantly modulated c-Fos density following acute (A) or chronic (B) LI-rTMS compared to the LI-rTMS-induced e-field. Left hemisphere: Simulated e-field in mV/mm induced by the LI-rTMS coil placed above lambda with a current of 1.83 mA/μs [17]. Right hemisphere: Top-down view of brain regions with significantly upregulated (yellow) or downregulated (pink) overall c-Fos density following acute (A) or chronic (B) LI-rTMS compared to sham controls. Videos showing the 3D model of significantly regulated regions can be found in supplementary materials (Movie S1-2), and for exploration in a 3D space, 3D objects for import into the Scalable Brain Atlas Composer [40] are included on our GitHub Repository.

Chronic stimulation induced significant density changes in fewer regions than acute stimulation, with mostly upregulation of c-Fos expression. The greatest difference in c-Fos activity was the upregulation of VMH, CEA, and VISC.

### Functional Connectivity Network

In altering brain-wide c-Fos expression, LI-rTMS also manipulated brain-wide functional network topology (Fig. 4a, b). These changes had minor effects on the density of statistically significant positively correlated activity in the network, with stimulation decreasing the density of such connections in acute conditions by a factor of 0.71 (0.17696 to 0.12560) and increasing this density in the chronic condition by a factor of 1.14 (0.105400 to 0.120200) (Fig. 4c). However, stimulation increased the density of statistically significant anti-correlations by factors of 4.38 in the acute condition (0.000457 to 0.00200) and 22.88 in the chronic condition (0.00153 to 0.003500; Fig. 4d). Together, these results suggest that LI-rTMS shifts brain-wide functional coactivation, coinciding with not statistically significant correlations becoming increasingly anti-correlated. These changes in correlation coefficient magnitude were most apparent across neuroanatomical regions within broader hierarchical groupings of the striatum, pallidum, and the amygdala.

**Figure 4.**
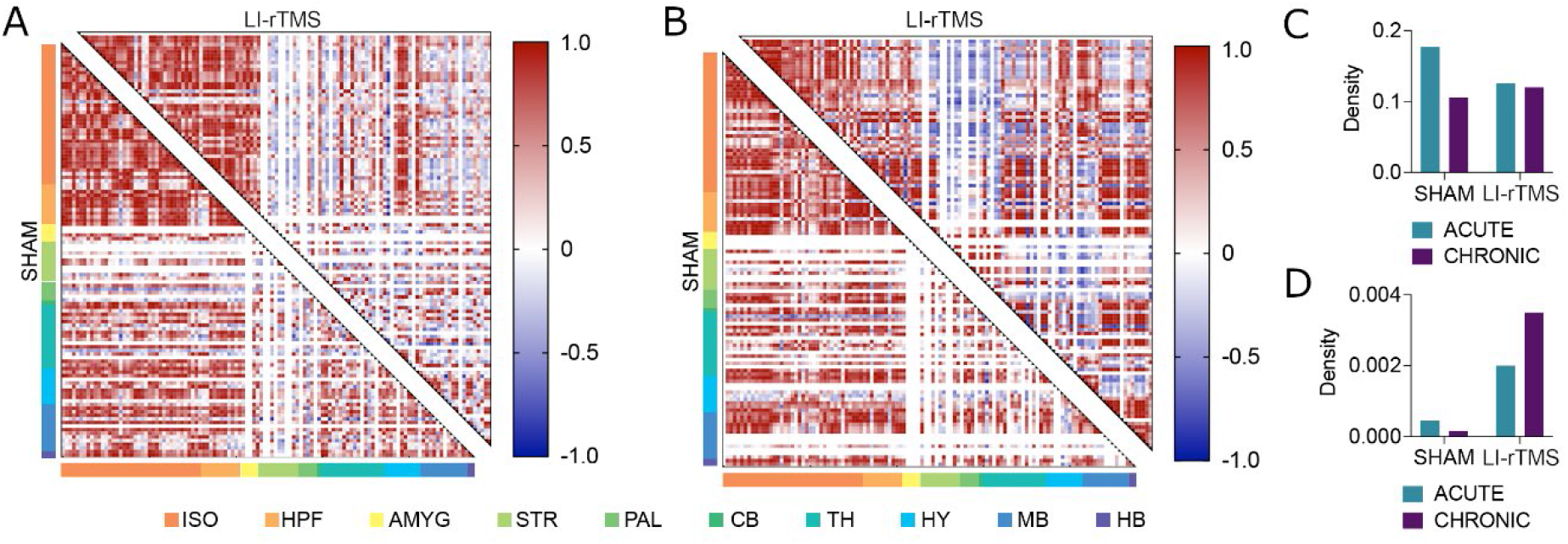
Brain-wide functional network topology and correlation density analyses. Functional network correlation matrices for sham (bottom left corner) and active LI-rTMS (top right corner) for acute **(A)** and chronic **(B)** stimulation groups. Matrices depict the coactivation of 115 neuroanatomical regions, with each row and column represent a single region and the intersection of rows and columns depicting the magnitude of the correlation between pairs of regions. Regions are also more broadly organised as isocortex (ISO), hippocampal formation (HPF), amygdala, (AMYG), striatum (STR), pallidum (PAL), cerebellum (CB), thalamus (TH), hypothalamus (HY), midbrain (MB), and hindbrain (HB). The prevalence of anticorrelations (depicted in blue) in the amygdala, pallidum, and striatum is increased considerably with LI-rTMS stimulation. Network density values, defined as the proportion of actual functional connections relative to the potential number of connections in a fully saturated network, show little change in **(C)** statistically significant positively correlated activity. **(D)** However, there was an increase in the density of anti-correlated activity in the network. See Supplementary File S1 for list of regions in the order than they are presented in the correlation matrices.

### Brain-Wide Parvalbumin and c-Fos Co-Expression

It has previously been demonstrated that LI-rTMS may influence parvalbumin interneurons underneath the coil e-field, as LI-rTMS can induce increases in cortical parvalbumin expression [25,41]. The altered brain-wide c-Fos expression patterns and network topology suggest that the effects of LI-rTMS extend beyond the site of the stimulation. To determine whether LI-rTMS affects activity of parvalbumin interneurons and whether changes also extended beyond the stimulation target, the brain-wide co-expression of parvalbumin and c-Fos was assessed. Representative images of parvalbumin and c-Fos co-expression are shown in Figure 5. The 2-way ANOVA for acute stimulation showed no significant effects or interaction. However, for chronic stimulation, there was a main effect of stimulation (F (1, 56) = 4.146, *p* = 0.0465), but no region effect or interaction (Fig 5a). Animals that received chronic stimulation showed significantly reduced % c-Fos^+^/PV^+^ cells. (Fig 5b). The effect of chronic stimulation, as reported by Hedge’s G, was most prevalent in the amygdala followed by the striatum (Fig 5c).

**Figure 5.**
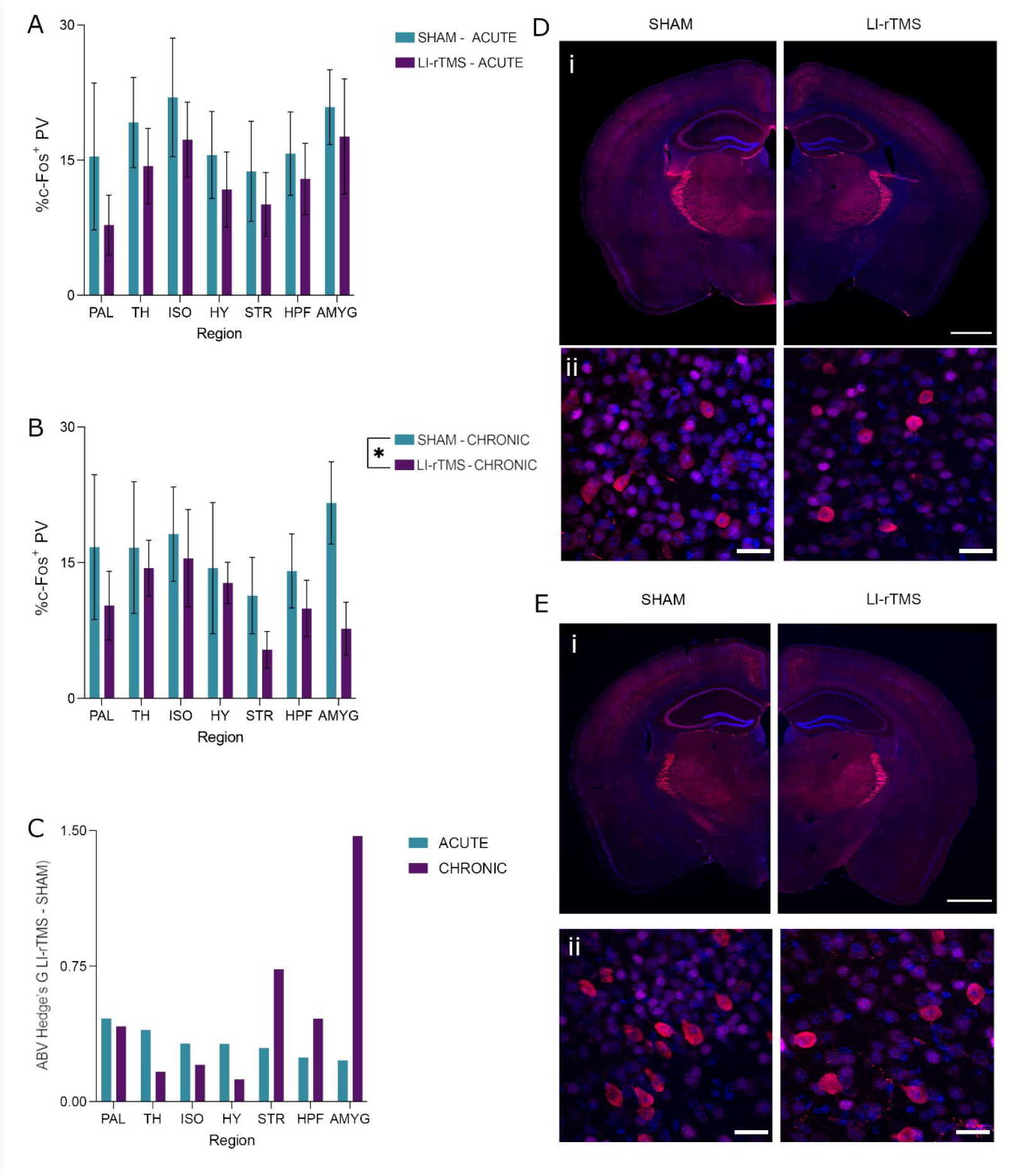
Percentage c-Fos/parvalbumin (PV) co-expression with acute or chronic LI-rTMS organised by brain region. **(A-B)** Percentage of parvalbumin cells co-expressing c-Fos in each region with acute (**A)** or chronic **(B)** LI-rTMS or sham controls. Error bars represent ±SEM **(C)** The absolute value (ABV) of Hedge’s G in both the acute and chronic groups. **(D-E)** Representational images of parvalbumin (red), c-Fos (magenta) and DAPI (blue) from the acute **(D)** and chronic **(E)** groups. Scale bars represent **(i)** 1000 µm or **(ii)** 25 µm.

## Discussion

Excitatory 10 Hz LI-rTMS caused widespread regulation of neuronal activity during stimulation. Both upregulation and downregulation of c-Fos expression occurred throughout the brain. The most prominent changes were during acute stimulation, particularly with upregulation of neuronal activity however, there were more limited activity changes with chronic stimulation. Changes to neuronal activity were present both underneath and away from the coil, suggesting direct and indirect induction of activity. LI-rTMS was also able to modulate functional connectivity on a brain-wide scale. LI-rTMS increased the extent to which regional c-Fos expression density was anti-correlated, with the most prominent changes occurring in the striatum, pallidum, and amygdala. This increase in anti-correlated activity was increasingly prominent with chronic stimulation. Potentially underlying the difference between acute and chronic stimulation effects, the activity of parvalbumin-positive interneurons across the brain decreased significantly with chronic LI-rTMS. These changes were most prominent in the striatum and amygdala, further corroborating the hypothesis that LI-rTMS manipulates parvalbumin interneuron activity to drive changes in functional connectivity. Overall, we show that LI-rTMS over the visual cortex appears to induce significant and widespread changes to the neuronal activity and functional connectivity of the brain, particularly in subcortical areas outside of the induced LI-rTMS e-field.

### LI-rTMS induces widespread changes to neuronal activity and functional connectivity

This study indicates that LI-rTMS modulates neuronal activity throughout the brain, but the effects of acute and chronic stimulation are subtly different. In the acute stimulation group, cortical regions directly beneath the coil showed prominent increases in activity compared to sham which suggests that electromagnetic induction using 10Hz LI-rTMS excites neurons within the induced e-field. While c-fos expression is an indirect marker of neuronal activity, our interpretation is supported by previous electrophysiological experiments showing that acute LI-rTMS lowers action potential thresholds *in vitro* in cortical neurons, resulting in increased neuronal excitability [42]. Interestingly, areas outside of the induced e-field also show significant regulation of neuronal activity, such as upregulation of c-Fos expression in striatal regions, and downregulation in several thalamic regions. These results support MRI experiments in rats demonstrating that the acute effects of LI-rTMS on neuronal activity extend beyond the site of stimulation [1,2], and are consistent with clinical neuroimaging studies of rTMS in humans [43,44]. LI-rTMS may thus acutely modulate interconnected regions *via* activation of downstream pathways.

Perhaps surprisingly, chronic LI-rTMS induced fewer changes to neuronal activity compared to acute LI-rTMS. However, chronic stimulation did result in more significant changes to brain-wide functional connectivity network topology, and these changes were most prevalent beyond the site of stimulation. The different outcomes of acute and chronic stimulation suggest that LI-rTMS effects are cumulative and may involve homeostatic mechanisms that prevent over-activation of neurons, as well as plasticity mechanisms that alter functional connectivity across brain regions. These processes are likely to underpin the long-term beneficial outcomes of therapeutic rTMS in depression and OCD and highlight the potential to optimise TMS treatment targets and protocols for specific dysfunctional networks.

### Chronic excitatory LI-rTMS modulates the activity of subcortical parvalbumin interneurons

The significance and origins of anti-correlations in c-Fos-based functional connectivity networks have been largely ignored [45–47]. Excitingly, the present study provides the first evidence for a link between parvalbumin interneurons and network anticorrelations. While previous studies have established that rTMS and LI-rTMS alter expression of parvalbumin [48,49], our study extends this work by showing that LI-rTMS significantly decreased the activity of parvalbumin interneurons. These largely GABAergic cell populations exert control over the activity of many more-abundant glutamatergic neuronal populations. Therefore, by altering the expression and activity of parvalbumin interneurons, LI-rTMS has the potential to modulate network synchronicity on a brain-wide scale [50–52]. This is in line with what we observed: regions in which parvalbumin interneuron activity was most prevalently modulated with chronic LI-rTMS (striatum and amygdala) coincide with the neuroanatomical regions in which neuronal c-Fos expression density became most prevalently anti-correlated. These results suggest that decreased parvalbumin interneuron activity promotes anti-correlated activity and is a potential mechanism through which LI-rTMS is able to modulate brain-wide functional connectivity.

The ability to modulate anticorrelated activity has numerous clinical implications, particularly through the lens of parvalbumin interneuron modulation. The magnitude of anticorrelated functional connectivity is dampened in several conditions, including depression [53,54], Parkinson’s disease [55], stroke [56], and anxiety [57]. These same conditions have also been demonstrated to have altered parvalbumin interneuron activity [58–61]. Many of the symptoms of these conditions have also been shown to improve with rTMS treatment [62–65]. Our results suggest that a possible mechanism through which LI-rTMS is able to ameliorate these symptoms is through its brain-wide modulation of parvalbumin interneuron activity and anticorrelated functional connectivity.

### Future Directions and Conclusion

Our research provides novel insight into how LI-rTMS changes functional connectivity at the cellular level, and forms part of a growing translational pipeline of preclinical and clinical neuromodulation studies that continue to inform human treatments. For example, our finding of changes in functional connectivity and parvalbumin interneuron activity in the amygdala and striatum provides new evidence that rTMS may be effective for treating disorders associated with aberrant activity in these regions. In addition, c-Fos density changes with acute LI-rTMS demonstrate that even short-term exposure to low-levels of electromagnetic fields can induce changes to neuronal activity throughout the brain, including in subcortical regions. While subthreshold rTMS effects remain poorly characterised in humans, low intensity magnetic fields are delivered as part of conventional, high-intensity rTMS, because magnetic field intensity decays with distance from the coil [18]. Therefore, our research showing changes with low-intensity fields is directly relevant to improving rTMS clinical outcomes in disorders characterised by dysregulation of subcortical circuitry.

## Acknowledgements

Funding for this study was provided in part by an NSERC Discovery Grant (RGPIN-2018-05135) to JRE. JM was supported by an Australian Government Research Training Program (RTP) scholarship, and Byron Kakulas Prestige scholarship. DJT received fellowships from NSERC and the Canadian Open Neuroscience Platform. JR was supported by a fellowship from Multiple Sclerosis Western Australia (MSWA). We acknowledge the Hotchkiss Brain Institute Advanced Microscopy Platform and the Cumming School of Medicine for support and use of the Olympus VS120-L100-W slide scanning microscope.

## Author Contributions

**Jessica Moretti:** Conceptualization, Methodology, Formal Analysis, Investigation, Visualisation, Writing – original draft, Writing – review & editing. **Dylan J. Terstege**: Methodology, Software, Formal Analysis, Investigation, Visualisation, Writing – original draft, Writing – review & editing. **Eugenia Z. Poh:** Methodology, Investigation, Writing – review & editing. **Jonathan R. Epp:** Writing – review & editing, Supervision, Resources. **Jennifer Rodger:** Writing – review & editing, Supervision, Resources.

## Supplementary figures

**Supplementary Figure S1.**
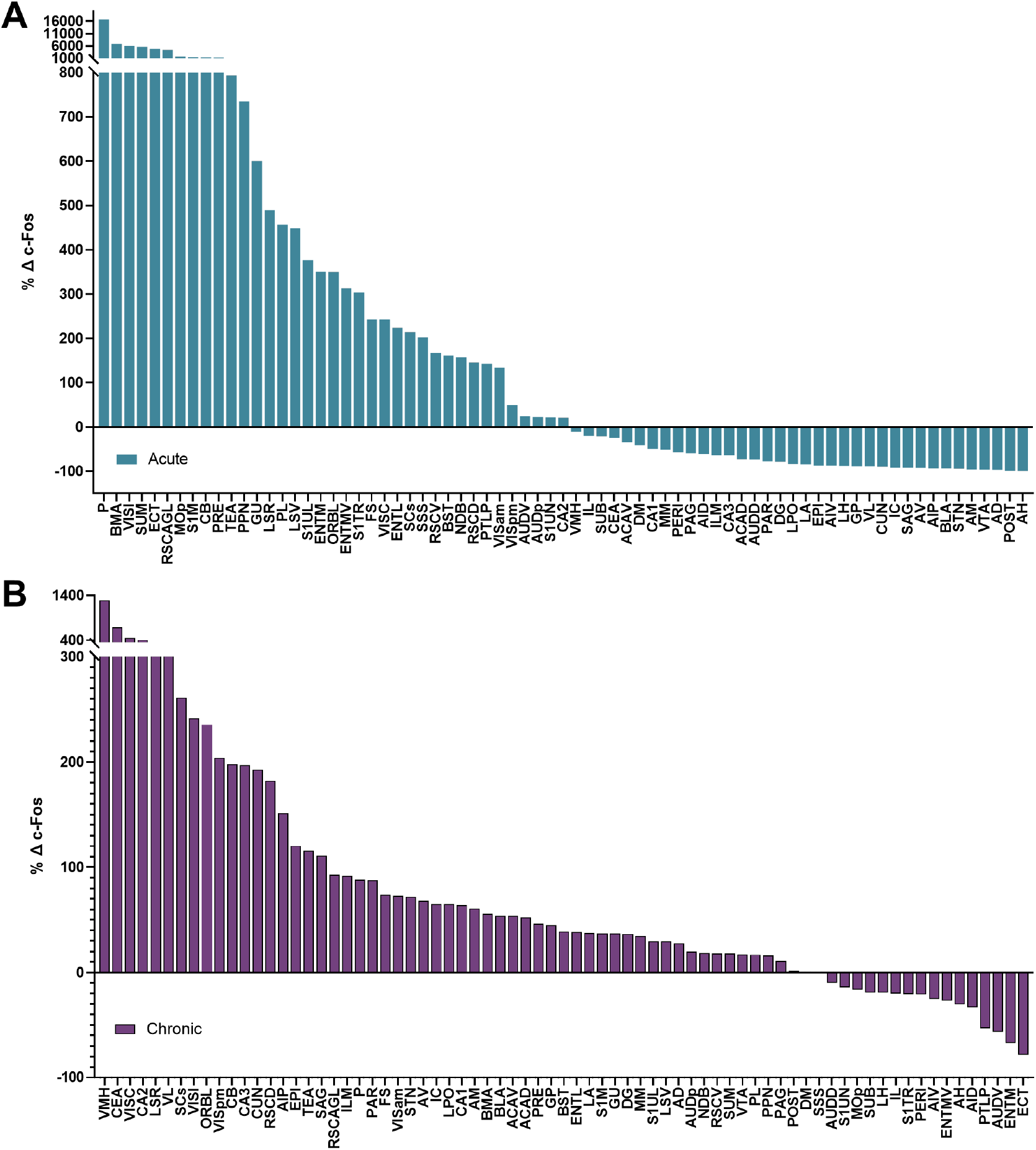
Mean percentage difference between active and sham LI-rTMS for (A) acute and (B) chronic groups for all analysed brain regions. Regions are organised by magnitude of percentage change. Note acute and chronic groups have different y-axis scales.

